# The next 20 years of genome research

**DOI:** 10.1101/020289

**Authors:** Michael C. Schatz

## Abstract

The last 20 years have been a remarkable era for biology and medicine. One of the most significant achievements has been the sequencing of the first human genomes, which has laid the foundation for profound insights into human genetics, the intricacies of regulation and development, and the forces of evolution. Incredibly, as we look into the future over the next 20 years, we see the very real potential for sequencing more than one billion genomes, bringing with it even deeper insights into human genetics as well as the genetics of millions of other species on the planet. Realizing this great potential, though, will only be achieved through the integration and development of highly scalable computational and quantitative approaches can keep pace with the rapid improvements to biotechnology. In this perspective, we aim to chart out these future technologies, anticipate the major themes of research, and call out the challenges ahead. One of the largest shifts will be in the training used to prepare the class of 2035 for their highly interdisciplinary world.

## I. Introduction

Modern quantitative biology is in many regards no different than previous eras, as biology has always benefited from quantitative analysis and modeling. The very foundation of modern genetics, Mendel’s principles of inheritance, was established through a quantitative analysis of some 30,000 pea plants and recognizing the inheritance of certain traits could be explained by a few simple mathematical rules [1]. This accomplishment well demonstrates the power of quantitative biology, especially considering that these laws were established in advance of modern molecular biology, including deciphering how genes encode for proteins, or that even DNA is the modality of inheritance.

At its core, modern quantitative biology follows the same principles as these early developments. It begins by asking questions of a biological system, and then recording and integrating observations made under varying conditions until some conclusion can be made. What is new in the modern era is the methods for collecting observations are now largely automated digital sensors, leading to much greater throughput and resolution. This includes the rise of DNA sequencing instruments, super-resolution digital microscopy, mass spectrometry, magnetic resonance imagery, or even satellite imagery used for studying biological systems.

While the instruments are now providing great quantities of data, they do not by themselves do any meaningful interpretation of them. Consequently, the second major advance in the field has been the increased importance of computational and analytical techniques used to study biological data. These approaches, including large-scale multi-core computing systems, advanced search and indexing algorithms, numerical optimization and modeling techniques, and many others, have primarily originated outside of biology in other quantitative disciplines. This new paradigm, often called “Biological Data Science” acknowledges that computer science, mathematics, physics, statistics, and other quantitative fields have developed advanced techniques that can be applied towards understanding biological data (https://datascience.nih.gov/).

The power of this approach comes from its ability to find relationships over very large numbers of observations, commonly stored in terabytes or petabytes of data [2]. Given the size and complexities of the relationships, though, this pursuit requires an end-to-end integration of approaches, forming an analysis stack starting with data collection and continuing through computational and statistical evaluations towards higher-level biological interpretations and insights **(Figure 1)**. Neglecting any layer of this stack will limit progress towards the higher goals: the instruments do not interpret data, the data will be inaccessible without high performance computing systems, and abstract quantitative approaches can be misled by technical artifacts or spurious correlations without deep understanding the underlying biology. Consequently, future advances in biology and medicine will come from the integration of biotechnologies, computational technologies, and quantitative reasoning, all designed by scientists with broad training.

**Figure 1.**
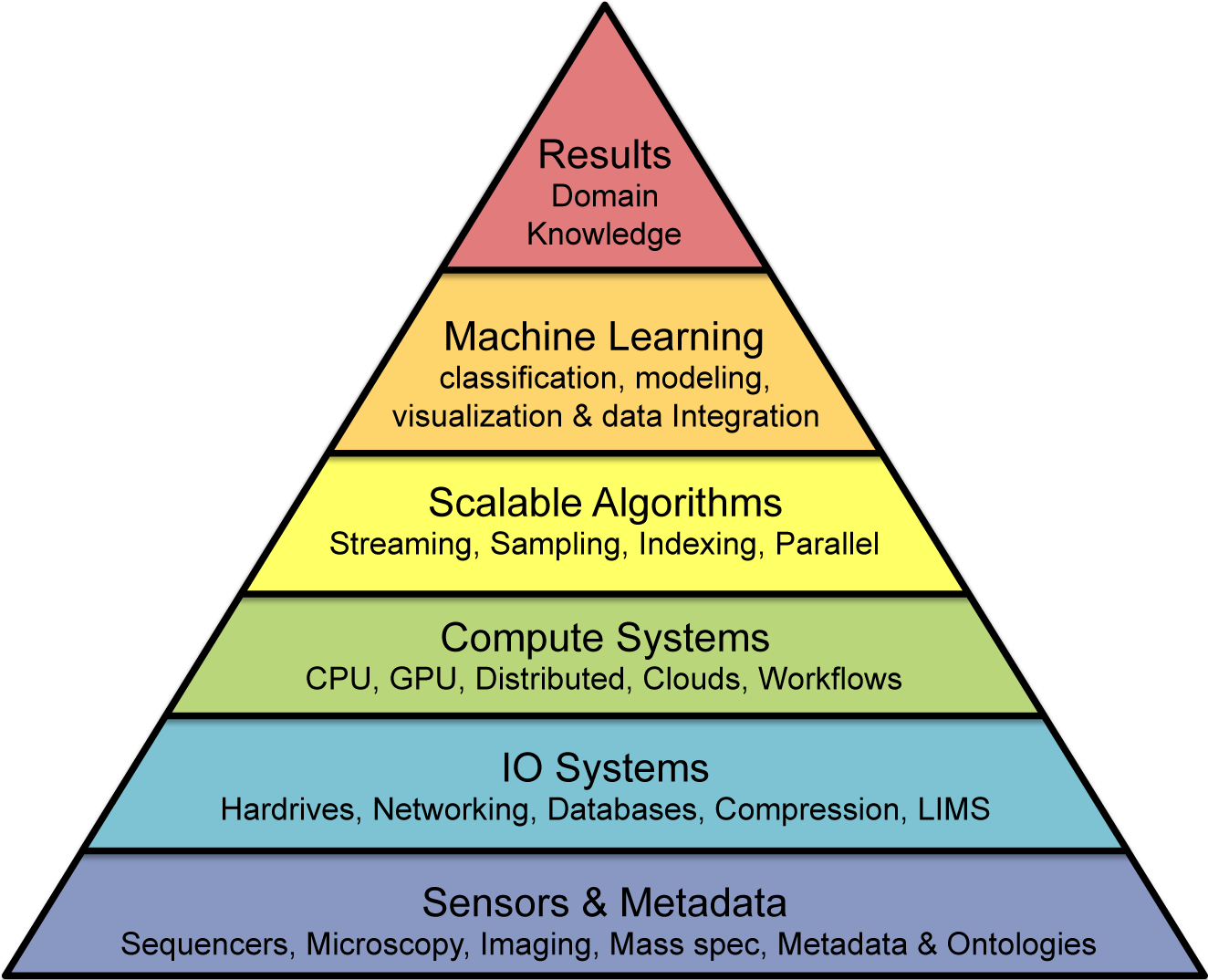
Quantitative Biology Analysis Stack. Large-scale projects in quantitative biology must address a multi-layer stack of approaches moving towards increasing levels of abstraction. At its base, the experiments begin with the technologies for collecting data and metadata from various biological sensors. The processing then proceed upwards through the IO and Compute layers that can support large-scale data processing, statistical and analysis software layers that can summarize and identifying trends in the data, until finally biological results can be achieved at the top leveraging the domain knowledge of the problem.

With this powerful combination of rapidly advancing biotechnology and rapidly advancing data science, quantitative biology has a very bright future over the next 20 years. Many fundamental questions in genomics and biology will start to be addressed regarding the structure and function of genomes, the molecular interplay within cells and organs, and the properties of entire species and ecosystems. One likely outcome from this research within human genetics will be an extended family tree linking together much of the world’s population, and with it unprecedented power to study the forces of inheritance as well as our own origins [3]. As we become more capable at interpreting genomes and monitoring molecular changes, we will also see profound advances in the medical community towards recognizing genetic risk factors and treating diseases based on one’s personal genomic makeup [4]. Outside of human genetics, we will see major efforts to use quantitative biology to enhance agriculture [5], monitor the microbiome [6], or understand the brain [7], among many other projects.

In this perspective we aim to chart out the major advances and challenges in quantitative biology over the next 20 years. This includes an analysis of what new biotechnologies and analytical tools are expected to develop, along with how those developments will lead to advances in biology and medicine. We end with a look into what technological and society challenges remain ahead, especially the shift in education that is needed to support the next generation of quantitative biologists.

## II. Advances in biotechnology

Quantitative biology is becoming an increasingly data-rich discipline, especially in the fields of genomics, systems biology, and computational neuroscience. Each of these fields has benefitted tremendously from recent improvements to sensor technology to probe into the inner workings of cells or environments with increasingly greater resolution. Interestingly, many of these advances have benefited from the core improvements to CCD technology leading to higher resolution and more affordable digital photography. These have naturally translated into improved microscopy or cytometry, but have had far-reaching benefits in other areas. For example, within DNA sequencing, improved CCD technology has enabled sequencing greater numbers and densities of molecules with fluorescently tagged nucleotides [8].

A premier example of how improved biotechnology has accelerated biology has been the development of high-throughput DNA sequencing. Originally developed in the 1970s, the initial protocols for sequencing DNA were slow and laborious with hazardous reagents [9]. These protocols were revolutionary at the time, but could only reliably sequence a few hundred or a few thousand base pairs per week per person, with substantial labor and reagent costs involved. In the past 20 years, though, several fully automated digital instruments have become commercially available that have dramatically accelerated the pace of sequencing [10]. During this time the worldwide capacity for sequencing DNA has doubled approximately every 9 to 12 months, and remarkably, the trillion fold improvements to throughput have been matched by similar reductions in costs. Today, the most powerful sequencing instrument available, the Illumina X10, can sequence the equivalent of one human genome every minute (3 Gbp / min) and has capacity to sequence 18,000 whole human genomes per year to deep coverage for about $1000 each [11].

Furthermore, the technologies used for sequencing DNA have been cleverly repurposed for several other non-DNA sequencing applications. This includes protocols for measuring the levels of mRNA transcription or translation inside of cells (RNA-seq, Ribo-seq), the presence of methylation (Methyl-seq), or the location and frequency of proteins binding to DNA (ChIP-seq), among dozens of other “omics” assays [12]. There have also been significant developments to push these technologies into ever more minute samples, including techniques to measure the genomes, transcriptomes, and epigenomes of individual cells, especially to probe the heterogeneity of gametes, cancer, the brain, and other complex samples [13].

Consequently genomics is now a data-rich and diverse subfield within modern quantitative biology, with studies exploring nearly every branch of the tree of life. There are sequencing instruments in more than 60 different countries on nearly every continent [14], and the worldwide sequencing capacity currently exceeds 35 petabases a year (35 million billion bases a year), enough capacity to sequence approximately 250,000 human genomes per year [15]. Much of that capacity is concentrated at research institutions, hospitals, and agricultural companies, and is used to study the genetics of humans and other species, especially those with medical, agricultural, or bioenergy importance. The sequencing company Illumina projects sequencing capacities will continue to double year over year, and projects by the end of 2017 more than 1.6 million human genomes will be sequenced [15]. Remarkably, at that rate of growth over the next 20 years the worldwide sequencing capacity will grow to reach more than 1 billion human genomes per year [16].

It is most likely that human and medical genomics, along with genomics of agriculture and energy production, will come to dominate the sequencing field as they have the largest economic incentives. In addition to widespread DNA sequencing, the various ‘omics assays will become increasingly important for monitoring health and disease over time [17]. This will be used, for example, to monitor diseases and recommend treatments based on changes to gene expression patterns before any macroscopic symptoms can be seen [18]. Single-cell approaches will also become increasingly important, especially for profiling circulating tumor cells in the blood before any recognizable tumors have developed [13]. Electronic medical records and personalized activity monitors, including Fitbits and mobile phones, will be used to continuously refine and update our understanding of our health and behavior [19].

Outside of medical genomics, many novel applications will develop as the instrumentation becomes smaller and less expensive [20]. One of the most important applications is the formation of a large distributed network of sequencers that can monitor for potential pathogens around the world. This sensor network will form the basis for a “digital immune system”, analogous to the worldwide weather network that can recognize changes to the composition of the viruses, microbes, and other agents in the environment [21]. Already there are biological sensors at high profile public sites, such as major sports arenas and transportation centers, used to monitor for pathogens passing in the air. Just as weather prediction becomes more informed and more accurate by worldwide monitoring, establishing a worldwide sequencing net would help us to monitor and contain epidemics before they can spread to larger populations.

Complementary to the technologies for measuring biological systems, new technologies for manipulating cells and synthesizing molecules will also become extremely important. Existing techniques for manipulating DNA or expression levels, such as restriction digests or RNAi [22], will be enhanced with new higher precision and more flexible technologies. Already, CRISPR/Cas9 systems are emerging as powerful techniques for editing genomes [23], and optogenetics can be used for real-time activation or repression of targeted cells [24]. Complementary advances in synthetic genomics are starting to yield entirely artificial genomes with genes to produce compounds of our own design [25].

Altogether, the next twenty years holds enormous potential for sensing and manipulating cells and molecules, creating an efficient feedback cycle between reading a genome or molecular activity, modeling its function, and measuring the effect of changing it.

## III. Computational and Quantitative Advances

Most immediately impacted by the massive growth to sequencing and senor technologies will be the computational systems used for storing and transferring biological data. For more than twenty years, NCBI and its international counterparts at the EBI and DDBJ have served as the central clearinghouse for genomic data [26]. Over the next twenty years, these resources will continue to steadily grow, although as the sequencing facilities grow from petabyte to exabyte scale, it will become less and less practical to transfer data into these archives as they exist today. Furthermore, as sequencing shifts from research purposes and into more direct medical applications, the incentive for making the data publically available in a centralized archive will be reduced or perhaps even legally restricted.

In its place, we will see the rise of federated approaches for exchanging biological data, especially computing centers dedicated to large sequencing facilities. Already this trend is beginning, and the NCBI Sequence Read Archive (SRA) currently only stores ∼1/10^th^ of the worldwide sequence production, around 3.8 Pbp of the more than 35Pbp sequenced so far (http://www.ncbi.nlm.nih.gov/Traces/sra/). Fortunately the rest of the data are not completely lost, and we are beginning to see the emergence of new exchange systems outside of traditional archives. These systems often consolidate regional and/or topical interests inside a dedicated cloud-based portal, such as CGHub [27] or ICGC [28] for consolidating cancer genomic data, or the recently launched BGI-cloud to provide access to the great resources available there (http://bgiamericas.com/data-analysis/bgi-cloud/). Illumina BaseSpace (https://basespace.illumina.com), DNAnexus (https://www.dnanexus.com/), Google Genomics (https://cloud.google.com/genomics/), and other commercial vendors are also emerging to help manage the deluge of data using commercial cloud platforms.

The major technical reason this model will become more widespread is that at large scales it is increasingly more efficient to upload code segments, often measured in kilobytes to megabytes, rather than to download entire large collections [29]. There are also economies of scale made possible through consolidating purchasing and time-sharing of equipment, especially for the many thousands of cores and petabytes of storage that will be need to be purchased and maintained. It also allows for the different cloud resources to specialize their services for different biological systems or political requirements: despite using common sequencing technologies, the tools and datasets needed for studying cancer are very different than those needed for studying crop development, as are the legal requirements in some countries compared to others.

While this model of federated cloud-based data warehousing offers many advantages, it also presents significant new challenges. Foremost, genomic data are most useful when they can be aggregated and combined in very large numbers. Otherwise, subtle genetic signals may be lost if the data are partitioned or if the measurements are recorded with incompatible formats. Therefore, the resources will have to establish common application programming interfaces (APIs) to enable remote access to their data, along with strong encryption and authentication safeguards to protect from theft or abuse. Today, the Global Alliance for Genomics and Health is one of the leading efforts to define such standards for genomics and other personal biomedical data, although even the most basic of federated tasks, so-called “Beacons” that identify if a resource has any individual with a particular mutation, are proving to be difficult to implement for mostly non-technical reasons (http://genomicsandhealth.org/our-work/current-initiatives/beacon-project).

Within each data warehouse, there will be major systems engineering challenges to consider: instead of a program crashing on one server, a code failure could disrupt thousands of cores at once leading to years or centuries of wasted computing effort. Most of the currently available genomics applications are not designed for this level of parallelism, needing new higher-level and easy to use workflow systems to orchestrate the scheduling and management of resources [30], as well as improved software engineering practices to build more robust software [31].

The massive data warehouses must also be built with tiered storage systems to prioritize access to the data within while keeping storage costs manageable [32]. Summary statistics, variant lists, expression profiles, and other highly processed data can be kept in active memory, while raw reads and other measurements can be archived to slower and less expensive medium. Specialized data compression algorithms [33], while extremely important to make the best use of every available byte, are unlikely to completely solve the storage needs. In an effort to control costs and complexity, there will be growing urgency to delete data as soon as possible. This will mark a radical shift in how data are currently viewed, and requires careful consideration of the “preciousness” of a sample: A research project exploring a rare cancer or ancient DNA sample will likely chose to archive everything, while studies of more abundant samples may elect to only store the processed results.

The applications and algorithms built for these warehouses will be focused on integrating the analysis over very large populations. Certain topics that are widely studied today, such as short read mapping or de novo assembly of individual genomes, will fade away as the algorithms mature and new sequencing technologies producing longer reads take over leading to extremely high quality genomes [34]. In its place will be the need for algorithms and systems for studying the genomes, transcriptomes, and epigenomes of millions of individuals at once, especially systems that can do so within a graph of sequence variability [35]. As sequencing becomes more widely used for real-time applications, such as a real-time readout of the transcription levels in a blood sample, the interpretation will need to be done on the fly without the raw data recorded at all. Already several streaming methods are becoming available for inferring expression levels from RNA-seq reads that are just as effective as their more traditional counterparts [36], and other omics data types inferring quantification from sequences are likely to follow similar developments.

Altogether, over the next twenty years we will see the development of major institutional and governmental data warehousing for millions to billions of genomes and other biomedical data. It will require maturing of the algorithms and formats used today into more scalable and interoperable systems as well as designing new systems to solve problems that are not even considered today. A growing need will be for streaming algorithms and other approaches that can make inferences over diverse data types as soon as they are produced, so that data storage needs will be as limited as possible.

## IV. Insights of Genome Research

If one of the greatest accomplishments from the last twenty years has been sequencing the first human genomes, one of the greatest pursuits for the next twenty years will be trying to understand what it all means. While we have been quite successful deciphering the genetic code of how genes code for proteins, the grander challenge of how genome sequences ultimately code for traits has remained largely a mystery. Answering this question is one of the central questions of quantitative biology and one of the great unanswered questions in all of science.

The rational for sequencing one million or one billion genomes using this powerful stack of technologies is not to produce one million or one billion separate lists of variants. Rather the hope is that the whole will be greater than the sum of its parts, and something new will emerge that sheds light on to what those sequences mean. Like Mendel, the hope is bringing together these data will lead to discovering the underlying patterns and rules of genetics and organism biology: who has which variants, how they are inherited or evolve, and how any variants are related to diseases and other traits. The insights we will gain will come in phases depending on many factors, including how many samples are available, what other measurements are available, and crucially, how complex the traits are under consideration. In some cases the relationships will be very clear, allowing us to leap almost immediately from genetic variants to their associated traits [37]. More often than not, though, this analysis will require a series of stepping stones to connect how a variant can alter the sequence or expression of a single gene, which in turn influences a pathway of interconnected genes, which then influences the overall cell or individual, all within in a particular environment [38].

Considering the multitude of environmental factors, cell-type specificities, and diversity of genetic backgrounds, understanding what each base of a genome fully means may be a never-ending pursuit. Nevertheless, over the next twenty years great strides will be made in raising our level of understanding. Today, we are most successful at interpreting major alterations within gene sequences [39], while we are substantially less informed about interpreting the relative importance and mechanisms for non-coding mutations [40]. Over the next twenty years, though, our power for doing so will greatly improve building on the pioneering work of ENCODE [41], the Roadmap Epigenomics Project [42], and similar projects that are starting to provide detailed annotations as to the roles and evolution of sequences all throughout the genome. The community is also currently largely focused on single nucleotide and other small variants, but as the sequencing technologies improve it is likely that the widespread nature and significance of structural variations will become even more pronounced [43]. Indeed, while a typical person has more SNPs than any other class of mutations, the total number of bases that are mutated, and hence the largest aspect of genetic diversity in a person, is mostly due to copy number and other structural variations [44, 45]. Finally, the widespread deployment of diverse sensors will allow us to more carefully consider environmental factors, such as Fitbit-type data of location and activities integrated with detailed molecular and microbiome profiles.

While extremely powerful, care must be taken when using these data and techniques to avoid unfounded generalizations or mistaking spurious correlations as casual relationships. Quantitative Biology needs integration of quantitative expertise with domain expertise to design the proper experiments, to fight technical artifacts, and to recognize relationships outside of the expected. As recently highlighted, while much scrutiny has been given to the importance of p-values, these evaluations come at the top of a complex analysis stack that requires careful scrutiny at every stage [46]. Whenever the result is most surprising or unexpected, that is exactly when one should be most critical of the methodology. Caution is also needed to consider the social changes and privacy risks that will come from building these massive data warehouses of personal information. It is surprisingly easy to identify someone from their genome today, and will become even easier and more accurate as these massive collections are established [47]. As genome sequences are connected to other personal data, the threat of genetic discrimination and abuse becomes real and ever lasting.

Nevertheless, the potential benefits for building and mining these massive collections are overall much larger than the potential risks. One of the major lessons learned from machine learning has been, when coupled with the right algorithms and systems, increasing the amount of data often leads to greatly increased performance and interpretation, including chess programs [48], translation systems [49], and even virtual Jeopardy contestants [50] that can outperform any human. Within genomics and quantitative biology, we hope and anticipate this will prove true as well, leading to many breakthroughs of profound significance over the next twenty years.

## VI. Recommendations for the Class of 2035

Children born this summer will grow up in a drastically different world than the childhoods of the current graduating class or those from twenty years ago. The class of 2035 will have unprecedented access to information, quantitative techniques, and biotechnologies that will be used to manipulate biological systems in currently unimaginable detail. While the foundations of biology will continue to be observation, experimentation, and interpretation, the technologies and approaches used will become ever more powerful and quantitative. More so than ever, we need to revise the curriculum to integrate computational and quantitative analysis as early as possible into their training so they are ready for the world ahead [51].

My recommendation to the class of 2035 is to embrace the integration of fields that is forming modern biology. To be competitive, you will need to establish a broad interdisciplinary foundation of math and sciences as well as strong communication skills. One of the most important skills you can develop early is computer programming. While sequencing technologies and other instrumentation will come and go over the next 20 years, biology will only continue to grow its dependency on computational analysis. And much like learning to speak a new language is often easier the younger you begin, learning to “speak” to a computer seems to follow a similar path. But I also encourage you to experiment with the “wet” side of biology as early as possible as well, since this will help you to appreciate the data you work with and put you in a position to run your own experiments end to end. Indeed, the most profound advances often occur at the intersection of new biotechnology and new quantitative analysis, when you can be the first to generate a novel data type that is used to unravel a mystery of how the world operates. Finally, always remember to keep focused on the most important problems that you can hope to address.

## Acknowledgements

I would like to thank all of the participants of the Biological Data Sciences conference, co-organized with Anne Carpenter and Matt Wood, held last fall at Cold Spring Harbor Laboratory that was inspirational to this work. I would also like to thank all of my great colleagues, friends, and family that have contributed to the ideas presented here. The project was supported in part by National Science Foundation award DBI-1350041.

## References

1. Mendel, G., Versuche über Plflanzen-hybriden. Verhandlungen des naturforschenden Vereines in Brünn, Bd. IV für das Jahr, 1865: p. 3–47.

2. Marx, V., Biology: The big challenges of big data. Nature, 2013. 498(7453): p. 255–60.

3. Ledford, H., Genome hacker uncovers largest-ever family tree. Nature, 2013.

4. Collins, F.S. and H. Varmus, A new initiative on precision medicine. N Engl J Med, 2015. 372(9): p. 793–5.

5. McCouch, S., et al., Agriculture: Feeding the future. Nature, 2013. 499(7456): p. 23–4.

6. Structure, function and diversity of the healthy human microbiome. Nature, 2012. 486(7402): p. 207–14.

7. Insel, T.R., S.C. Landis, and F.S. Collins, Research priorities. The NIH BRAIN Initiative. Science, 2013. 340(6133): p. 687–8.

8. Ansorge, W.J., Next-generation DNA sequencing techniques. New Biotechnology, 2009. 25(4): p. 195–203.

9. Sanger, F. and A.R. Coulson, A rapid method for determining sequences in DNA by primed synthesis with DNA polymerase. J Mol Biol, 1975. 94(3): p. 441–8.

10. Reuter, J.A., D.V. Spacek, and M.P. Snyder, High-Throughput Sequencing Technologies. Mol Cell, 2015. 58(4): p. 586–597.

11. Illumina. HiSeq X Series of Sequencing Systems. 2015 [cited 2015 April 28, 2015]; Available from: http://www.illumina.com/content/dam/illumina-marketing/documents/products/datasheets/datasheet-hiseq-x-ten.pdf.

12. Soon, W.W., M. Hariharan, and M.P. Snyder, High-throughput sequencing for biology and medicine. Molecular systems biology, 2013. 9: p. 640.

13. Wigler, M., Broad applications of single-cell nucleic acid analysis in biomedical research. Genome medicine, 2012. 4(10): p. 79.

14. Omicsmap. Next Generation Genomics: World Map of High-throughput Sequencers. 2015; Available from: http://omicsmaps.com/.

15. Regalado, A. EmTech: Illumina Says 228,000 Human Genomes Will Be Sequenced This Year. MIT Technology Review 2014 [cited 2015 April 28, 2015]; Available from: http://www.technologyreview.com/news/531091/emtech-illumina-says-228000-human-genomes-will-be-sequenced-this-year/.

16. Stephens, Z.D., et al., Big Data: Astronomical or Genomical? PLoS Biology, 2015. In Press.

17. Chen, R., et al., Personal omics profiling reveals dynamic molecular and medical phenotypes. Cell, 2012. 148(6): p. 1293–307.

18. Simon, R. and S. Roychowdhury, Implementing personalized cancer genomics in clinical trials. Nat Rev Drug Discov, 2013. 12(5): p. 358–69.

19. Gottesman, O., et al., The Electronic Medical Records and Genomics (eMERGE) Network: past, present, and future. Genet Med, 2013. 15(10): p. 761–71.

20. Rusk, N., MinION takes center stage. Nat Methods, 2015. 12(1): p. 12–3.

21. Schatz, M.C. and A.M. Phillippy, The rise of a digital immune system. GigaScience, 2012. 1(1): p. 4.

22. Hannon, G.J., RNA interference. Nature, 2002. 418(6894): p. 244–51.

23. Jinek, M., et al., A programmable dual-RNA-guided DNA endonuclease in adaptive bacterial immunity. Science, 2012. 337(6096): p. 816–21.

24. Deisseroth, K., Optogenetics. Nat Methods, 2011. 8(1): p. 26–9.

25. Gibson, D.G., et al., Creation of a bacterial cell controlled by a chemically synthesized genome. Science, 2010. 329(5987): p. 52–6.

26. Coordinators, N.R., Database resources of the National Center for Biotechnology Information. Nucleic Acids Res, 2015. 43(Database issue): p. D6–17.

27. Wilks, C., et al., The Cancer Genomics Hub (CGHub): overcoming cancer through the power of torrential data. Database (Oxford), 2014. 2014.

28. International Cancer Genome C., et al., International network of cancer genome projects. Nature, 2010. 464(7291): p. 993–8.

29. Schatz, M.C., B. Langmead, and S.L. Salzberg, Cloud computing and the DNA data race. Nature biotechnology, 2010. 28(7): p. 691–3.

30. Afgan, E., et al., Harnessing cloud computing with Galaxy Cloud. Nat Biotechnol, 2011. 29(11): p. 972–4.

31. Wilson, G., et al., Best practices for scientific computing. PLoS Biol, 2014. 12(1): p. e1001745.

32. Haussler, D., et al., A Million Cancer Genome Warehouse. 2012, EECS Department, University of California, Berkeley.

33. Hsi-Yang FritzM., et al., Efficient storage of high throughput DNA sequencing data using reference-based compression. Genome research, 2011. 21(5): p. 734–40.

34. Berlin, K., et al., Assembling large genomes with single-molecule sequencing and locality-sensitive hashing. Nat Biotechnol, 2015.

35. Church, D.M., et al., Extending reference assembly models. Genome Biol, 2015. 16: p. 13.

36. Patro, R., S.M. Mount, and C. Kingsford, Sailfish enables alignment-free isoform quantification from RNA-seq reads using lightweight algorithms. Nature biotechnology, 2014. 32(5): p. 462–4.

37. McCarthy, M.I., et al., Genome-wide association studies for complex traits: consensus, uncertainty and challenges. Nat Rev Genet, 2008. 9(5): p. 356–69.

38. Wang, K., M. Li, and H. Hakonarson, Analysing biological pathways in genome-wide association studies. Nat Rev Genet, 2010. 11(12): p. 843–54.

39. Wang, K., M. Li, and H. Hakonarson, ANNOVAR: functional annotation of genetic variants from high-throughput sequencing data. Nucleic Acids Research, 2010. 38(16): p. e164.

40. Ward, L.D. and M. Kellis, Interpreting noncoding genetic variation in complex traits and human disease. Nat Biotechnol, 2012. 30(11): p. 1095–106.

41. Dunham, I., et al., An integrated encyclopedia of DNA elements in the human genome. Nature, 2012. 489(7414): p. 57–74.

42. Romanoski, C.E., et al., Epigenomics: Roadmap for regulation. Nature, 2015. 518(7539): p. 314–6.

43. Chaisson, M.J., et al., Resolving the complexity of the human genome using single-molecule sequencing. Nature, 2015. 517(7536): p. 608–11.

44. Sebat, J., et al., Large-scale copy number polymorphism in the human genome. Science, 2004. 305(5683): p. 525–8.

45. Zarrei, M., et al., A copy number variation map of the human genome. Nat Rev Genet, 2015. 16(3): p. 172–83.

46. Leek, J.T. and R.D. Peng, Statistics: P values are just the tip of the iceberg. Nature, 2015. 520(7549): p. 612.

47. Erlich, Y. and A. Narayanan, Routes for breaching and protecting genetic privacy. Nature reviews. Genetics, 2014. 15(6): p. 409–21.

48. Campbell, M., A.J. Hoane, and F.-h. Hsu, Deep blue. Artificial intelligence, 2002. 134(1): p. 57–83.

49. Koehn, P., et al., Moses: open source toolkit for statistical machine translation, in Proceedings of the 45th Annual Meeting of the ACL on Interactive Poster and Demonstration Sessions. 2007, Association for Computational Linguistics: Prague, Czech Republic. p. 177–180.

50. Ferrucci, D.A., Introduction to ”This is Watson”. IBM Journal of Research and Development, 2012. 56(3.4): p. 1:1–1:15.

51. Schatz, M.C., Computational thinking in the era of big data biology. Genome Biology, 2012. 13(11): p. 177.

